# IL-4 and helminth infection downregulate MINCLE-dependent macrophage response to mycobacteria and Th17 adjuvanticity

**DOI:** 10.1101/2021.08.17.456702

**Authors:** Judith Schick, Matthew Lacorcia, Meltem Altunay, Christoph Schubart, Barbara Bodendorfer, Dennis Christensen, Christian Alexander, Stefan Wirtz, David Vöhringer, Clarissa Prazeres da Costa, Roland Lang

**Affiliations:** Institut für Klinische Mikrobiologie, Immunologie und Hygiene, Universitätsklinikum Erlangen, Friedrich-Alexander Universität Erlangen-Nürnberg, Germany; Institut für Medizinische Mikrobiologie, Immunologie und Hygiene, Center for Global Health, Technische Universität München, Germany; Infektionsbiologische Abteilung, Universitätsklinikum Erlangen, Friedrich-Alexander Universität Erlangen-Nürnberg, Germany; Adjuvant Research, Dept. of Infectious Disease Immunology, Statens Serum Institut, Copenhagen, Denmark; Cellular Microbiology, Forschungszentrum Borstel, Leibniz Lung Center Borstel, Germany; Medizinische Klinik 1, Universitätsklinikum Erlangen, Friedrich-Alexander Universität Erlangen-Nürnberg, Germany

## Abstract

The myeloid C-type lectin receptor (CLR) MINCLE senses the mycobacterial cell wall component trehalose-6,6-dimycolate (TDM). Recently, we found that IL-4 down-regulates MINCLE expression in macrophages. IL-4 is a hallmark cytokine in helminth infections, which appear to increase the risk for mycobacterial infection and active tuberculosis. Here, we investigated functional consequences of IL-4 and helminth infection on MINCLE-driven macrophage activation and Th1/Th17 adjuvanticity. IL-4 inhibited MINCLE and cytokine induction after macrophage infection with *Mycobacterium bovis* Bacille Calmette-Guerin (BCG). Infection of mice with BCG upregulated MINCLE on myeloid cells, which was inhibited by IL-4 plasmid injection and by infection with the nematode *Nippostrongylus brasiliensis* in monocytes. To determine the impact of helminth infection on MINCLE-dependent immune responses, we vaccinated mice with a recombinant protein together with the MINCLE ligand Trehalose-6,6-dibehenate (TDB) as adjuvant. Concurrent infection with *N. brasiliensis* or with *Schistosoma mansoni* promoted T cell-derived IL-4 production and suppressed Th1/Th17 differentiation in the spleen. In contrast, helminth infection did not reduce Th1/Th17 induction by TDB in draining peripheral lymph nodes, where IL-4 levels were unaltered. Upon use of the TLR4-dependent adjuvant G3D6A, *N. brasiliensis* infection impaired selectively the induction of splenic antigen-specific Th1 but not of Th17 cells. Thus, helminth infection attenuated the Th17 response to MINCLE-dependent immunization in an organ-specific manner. Taken together, our results demonstrate down-regulation of MINCLE expression on monocytes and macrophages by IL-4 as a possible mechanism of thwarted Th17 vaccination responses by underlying helminth infection.

## Introduction

Tuberculosis kills more than 1 million humans each year and is easily spread by aerosol infection. Importantly, after exposure to *Mycobacterium tuberculosis* (MTB) by inhalation, only a minority of contacts develops active pulmonary disease with mycobacterial replication and destruction of lung tissue. In the majority of exposed people, inhaled MTB is either killed by the innate immune system of the lung, or triggers the development of granuloma structures around infected cells [1]. The development of specific CD4+ T cell responses then induces control of mycobacterial growth without achieving eradication of MTB, a status referred to as latency [1]. It is estimated that more than a quarter of the world’s population is latently infected with MTB. Their life-long risk to develop reactivation tuberculosis is around 5% and is influenced by a variety of genetic and environmental factors.

Parasitic helminth infections with intestinal nematodes, filaria or trematodes affect an estimated number of > 2 billion humans [2]. Similar to tuberculosis, helminth infections are often chronic and not acutely life-threatening. Epidemiologically, tuberculosis and helminth infections co-occur in many tropical and subtropical countries [3]. Immune responses to helminths in general are characterized by a strong Th2 component with production of IL-4, IL-5, IL-13, IL-10 and IgE [2], whereas protective immunity to MTB requires robust Th1 immunity and is possibly enhanced by Th17 cells [4, 5]. These diametrically opposed immunological biases and effector mechanisms raise the question how MTB and parasitic helminths mutually influence each other in co-infected individuals [2]. Interestingly, helminth infection of household contacts of patients with active, smear-positive tuberculosis increased the rate of tuberculin skin test conversion indicating a higher risk to become infected [6]. Furthermore, active tuberculosis patients are more often co-infected with helminths than healthy counterparts [7, 8]. Furthermore, helminth co-infection is associated with more advanced disease in tuberculosis patients and with reduced production of IFNγ but increased IL-10 [9]. These findings indicate that the type 2 immune bias in helminth-infected individuals can impair the development of protective adaptive immunity to MTB. In contrast, other studies showed that underlying helminth infection can promote a Th2 and regulatory T cell response to MTB with a reduced risk to develop open, smear-positive pulmonary tuberculosis [10]. In mice, infection with *Nippostrongylus brasiliensis* enhances bacterial load in the lung after MTB infection, an effect attributed to the action of alternatively activated macrophages [11].

Underlying helminth infection may also influence the response to vaccination with *M. bovis* Bacille Calmette-Guerin (BCG), which shows large geographic variation with reduced efficacy in countries with a high prevalence of worm infections [12, 13]. Experimental infection of mice with *Schistosoma (S*.*) mansoni* or *Heligmosomoides (H*.*) polygyrus* impaired Th1 cell generation after vaccination with BCG [14, 15], although another study found that chronic enteric infection with *H. polygyrus* did not interfere with primary or memory T cell responses to BCG [16]. Thus, these results from experimental mouse models show that not all helminth infections have a similarly strong impact on protective immunity to MTB. Anthelminthic treatment prior to BCG administration led to increased IFNγ and IL-12 production [17]. It is noteworthy that maternal helminth infection during pregnancy negatively affected the frequency of IFNγ-producing T cells in cord blood of neonates [18] and the development of Th1 immunity in BCG-vaccinated offspring [19]. However, attempts to enhance the immunogenicity of BCG vaccination by anthelminthic treatment of mothers during pregnancy were not successful [20].

Sensing of mycobacteria by the innate immune system is essential for a robust defense reaction, including the secretion of chemokines to attract leukocytes to the site of infection and the production of cytokines directing antigen-specific T cell responses towards Th1 and Th17 differentiation. Mycobacteria contain a plethora of PAMPs, e.g. the TLR ligands 19 kDa lipoprotein, lipoarabinomannan (LAM) and mycobacterial DNA rich in CpG motifs [21]. The hydrophobic cell wall of mycobacteria is especially rich in the glycolipid trehalose-6,6’-dimycolate (TDM, *aka* the cord factor), which is sufficient to trigger granulomatous reactions *in vivo* [22], induces inflammatory cytokine production by macrophages [23] and confers the Th1/Th17 adjuvant activity known from Freund’s Complete Adjuvant (CFA) [24]. TDM binds to the C-type lectin receptor (CLR) MINCLE (encoded by the *Clec4e* gene) [22, 25], which is inducibly expressed in macrophages [26] and associates with the adapter protein Fc receptor gamma chain (FcRγ, encoded by *Fcerg1*) [27]. Signaling by MINCLE requires binding of the kinase SYK to the phosphorylated ITAM of the FcRγ, and proceeds via the CARD9-BCL10-MALT1 complex to activation of NFκB and expression of inflammatory target genes [28, 29]. In addition to TDM, several additional glycolipids from the mycobacterial cell wall were recently identified as ligands for CLR, most of them like MINCLE members of the FcRγ-coupled DECTIN-2 family [30]. MCL, closely related to MINCLE and able to heterodimerize with it, also can bind TDM and in addition glycero-monomycolate [31, 32]; DECTIN-2 binds mannose-capped LAM [33]; and DCAR is a receptor for mycobacterial phosphatidyl-inositol mannosides [34]. The prototypic myeloid signaling CLR, DECTIN-1 directly recruits SYK through its non-classical ITAM [35]; it, too, binds and is activated by an yet unidentified mycobacterial ligand [36]. Since the signals emanating from the interaction of multiple CLR with MTB are transmitted through SYK-CARD9, it is not surprising that CARD9-deficient mice are highly susceptible to infection with MTB and succumb with high bacterial burden in the organs [37]. Knockout mice for individual CLR showed less severe phenotypes but were, dependent on the infection model employed, moderately more susceptible to infection [38-41].

Already in the first report on MINCLE, the inducible expression in response to TLR stimulation with LPS and by the cytokine TNF was reported [26]. Indeed, TNF is essential for the upregulation of MINCLE in macrophages stimulated with the cord factor and for the Th17 promoting effect of the adjuvant CAF01 that contains the MINCLE ligand TDB [42]. In contrast, the Th2 cytokines IL-4 and IL-13 downregulate expression of MINCLE, MCL and DECTIN-2, in human monocytes/macrophages and DC, as well as in mouse bone marrow-derived macrophages [43, 44]. This inhibitory effect of the Th2 cytokines IL-4 and IL-13 depends on the transcription factor STAT6 and is associated with a reduced production of G-CSF and TNF by macrophages stimulated with the MINCLE ligand TDB, but not after stimulation with LPS [43]. Thus, expression of the DECTIN-2 family CLR MINCLE, MCL and DECTIN-2 is subject to a substantial regulation by the cytokines TNF and IL-4, which are key factors in type 1 and type 2 immune responses, respectively. Since type 2 immunity is a hallmark of many helminth infections, the question arises whether the expression and function of DECTIN-2 family CLR may be compromised during helminth infection *in vivo* [45].

Here, we have investigated the functional impact of IL-4 and helminth infection on myeloid responses to mycobacteria or the cord factor-based adjuvant TDB *in vitro* and *in vivo*. Our data show that IL-4 impairs MINCLE expression on monocytes *in vivo*, but does not interfere with phagocytosis of BCG. Underlying helminth infection attenuated immune responses to recombinant protein together with different adjuvants in the spleen but not in draining lymph nodes. The specific impairment of Th17 responses to MINCLE-dependent adjuvant indicates the contribution of downregulation of DECTIN-2 family CLR to vaccine antagonism induced by helminth infection.

## Results

### IL-4 impairs upregulation of MINCLE and other DECTIN-2 family CLR in macrophages stimulated with BCG

In previous work, we observed that IL-4 downregulates expression of MINCLE, MCL and DECTIN-2 when macrophages were stimulated with the MINCLE-ligand TDB, but not after triggering TLR4 with LPS [43]. Since mycobacteria contain multiple ligands for CLR and TLR, we were interested whether IL-4 has any impact on changes in DECTIN-2 family CLR expression in macrophages stimulated with *M. bovis* BCG (Fig. 1). Stimulation of BMM with BCG caused the strongest induction of MINCLE (up to 400-fold after 48 hours) and was reduced significantly by IL-4 (Fig. 1A). A similar pattern of inducibility and regulation by IL-4 was observed for DECTIN-2 and MCL, albeit the increases at the mRNA level were more moderate (e.g. 50-fold for DECTIN-2 and 8-fold for MCL after stimulation with BCG) (Fig. 1B, C). IL-4 did not inhibit, but actually increased expression of DECTIN-1 (Fig. 1D), confirming a selective impact on DECTIN-2 family CLR. The inhibitory effect of IL-4 on BCG-induced expression of MINCLE was also evident at the protein level, when analysed by flow cytometry (Fig. 1E).

**Figure 1:**
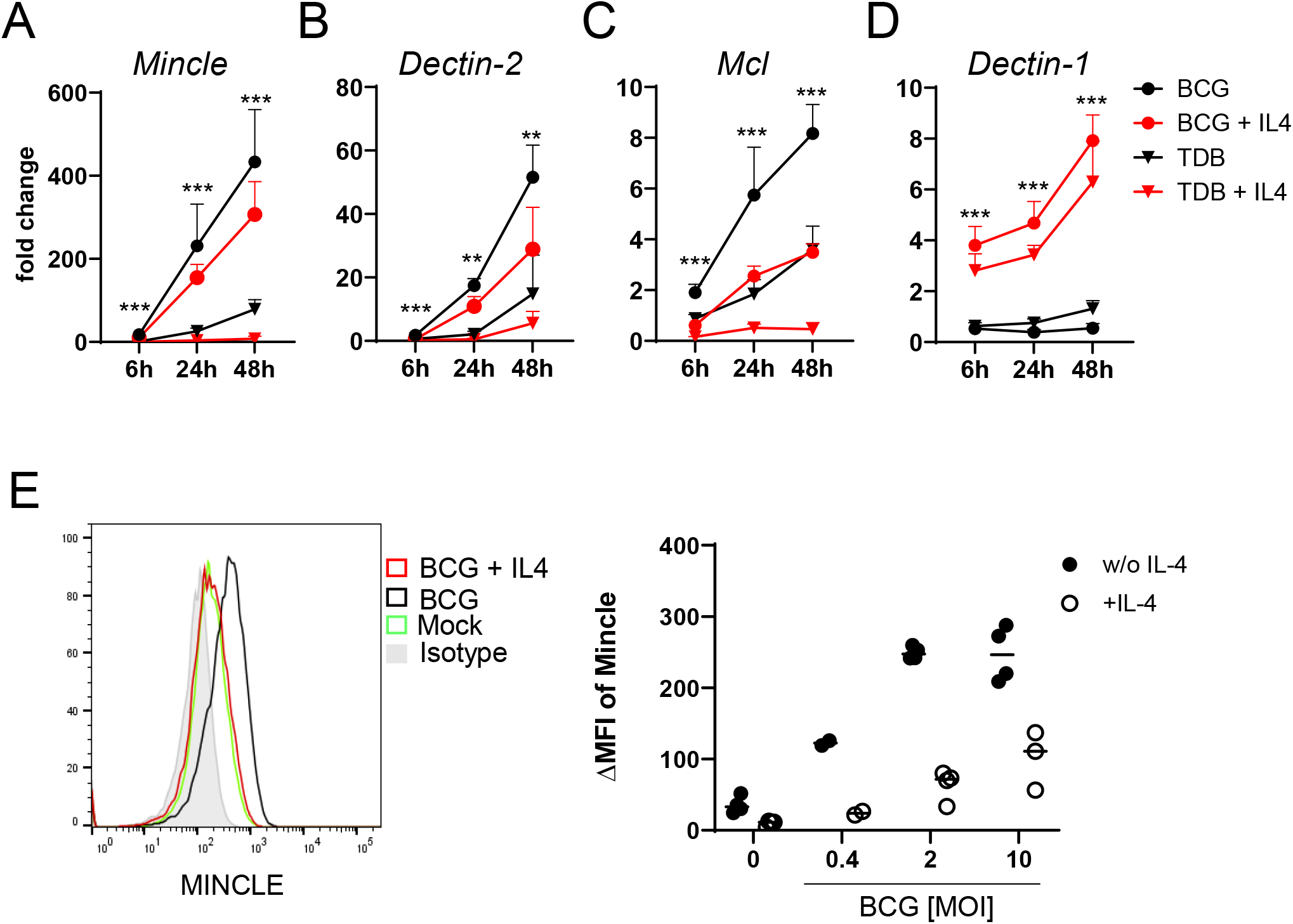
IL-4 impairs upregulation of MINCLE and other DECTIN-2 family CLR in macrophages stimulated with BCG. (A-D) C57BL/6 BMM were stimulated as indicated in presents or absence of IL-4 for 6, 24 or 48h. MINCLE (A), DECTIN-2 (B), MCL (C), and DECTIN-1 (D) mRNA expression was determined by qRT-PCR shown as fold change calibrated to unstimulated control. Data are depicted as mean + SD from two independent experiments performed in biological duplicates. (E) BMM were stimulated with BCG at the indicated MOI in the presence or absence of IL-4 for 24 hours, followed by staining of MINCLE expression. Representative stainings are shown as histogram overlay (left panel), for quantification (right panel) the median fluorescence intensity of the isotype control staining was subtracted from MINCLE signal to obtain the ΔMFI. Each point represents one mouse, pooled from two experiments. *p < 0.05, **p < 0.01, ***p < 0.001.

### IL-4 does not affect phagocytosis of BCG, but inhibits cytokine production

Several CLR, including DECTIN-2 and MCL, are phagocytic receptors [46, 47]. Therefore, we next measured phagocytosis of fluorescent BCG (expressing DsRed) by flow cytometry in the presence or absence of IL-4 (Fig. 2A). BMM infected with BCG-DsRed in different MOI dose-dependently ingested the mycobacteria, IL-4 had no impact on phagocytosis after 6 hours (not shown) and 24 hours (Fig. 2A). In contrast, the high levels of the cytokines G-CSF and TNF in supernatants of BMM infected with BCG were significantly reduced by co-treatment with IL-4 (Fig. 2B), extending our previous results for TDB [43] to the response induced by whole mycobacteria. Stimulation of BMM deficient in MINCLE or in FcRγ chain, the adapter protein used by MINCLE, MCL and DECTIN-2, with BCG led to a reduced production of TNF and G-CSF, confirming the contribution of this receptor to macrophage activation by mycobacteria [22, 25] (Fig. 2B). Of interest, the inhibitory effect of IL-4 was still visible in MINCLE-deficient BMM for G-CSF but not for TNF; in addition, it was much less pronounced than in WT BMM (Fig. 2B), indicating that downregulation of MINCLE is a prominent mechanism of IL-4-induced impairment of macrophage cytokine production to mycobacteria.

**Figure 2:**
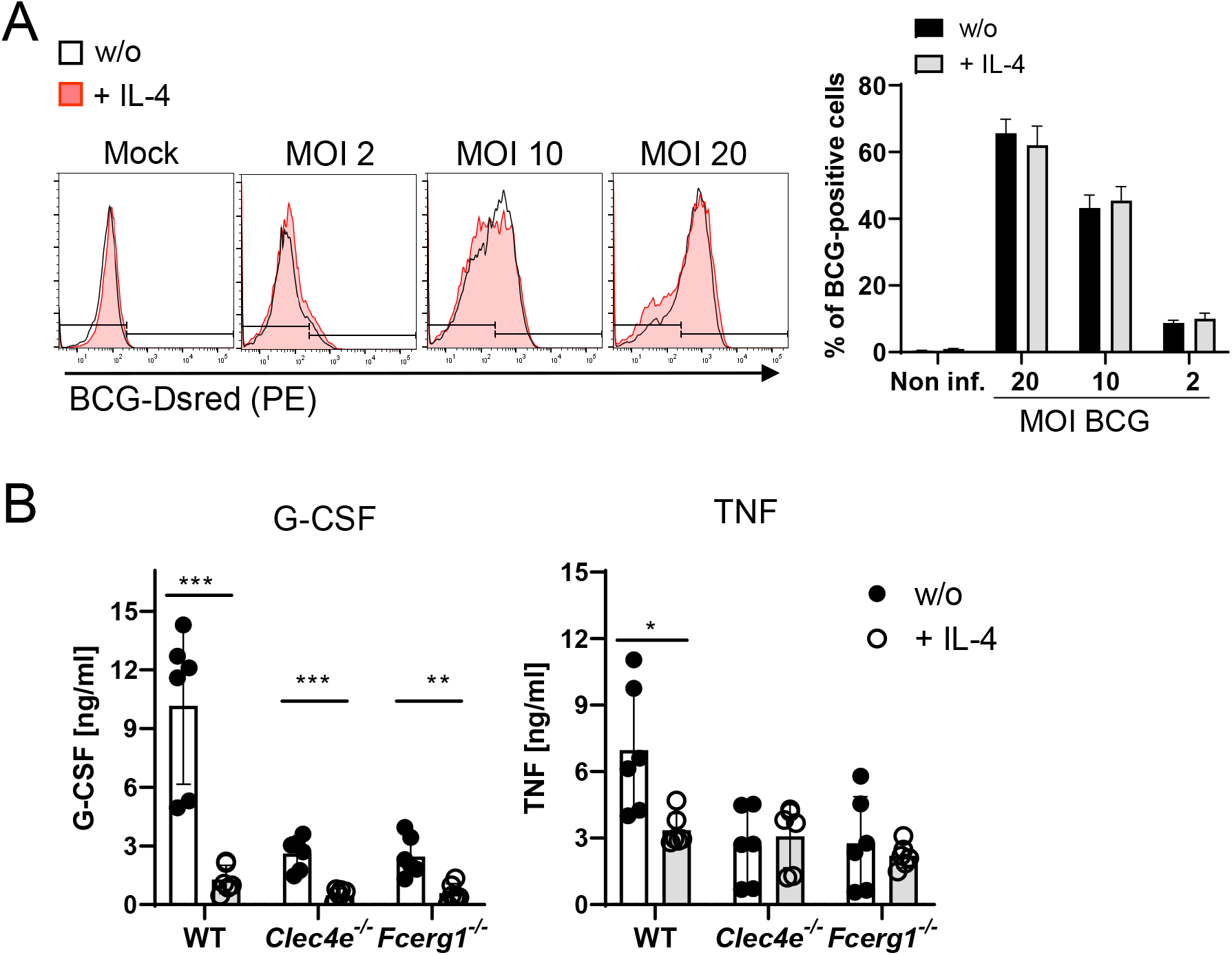
IL-4 does not affect phagocytosis of BCG but inhibits cytokine production. **(A)** C57BL/6 BMM were infected with different MOI of fluorescent BCG-Dsred co-treated with IL-4 or not as indicated. Phagocytic uptake was measured via flow cytometry. **(A)** Representative histograms show phagocytosis of BCG by detection of fluorescent DsRed signal in BMMs. Quantitative analysis of phagocytosis based on percentage of BCG-positive cells at 6 and 24 h post infection. Data is depicted from 2-3 independent experiments performed in biological duplicates. **(B)** C57BL/6 *(WT)*, MINCLE knockout *(Clec4e*^*-/-*^) and FcRγ (*Fcerg1*^*-/-*^) BMM were stimulated with BCG for 24 h. Production of G-CSF and TNF was measured from cell culture supernatants via ELISA. Data is depicted from 3 independent experiments performed in biological duplicates. *p < 0.05, **p < 0.01, ***p < 0.001.

### Overexpression of IL-4 inhibits MINCLE expression after intraperitoneal infection with BCG

Given the impact of IL-4 on BCG-induced MINCLE expression *in vitro*, we next used intraperitoneal infection of mice with BCG for analysis of MINCLE regulation by IL-4 *in vivo*. Hydrodynamic injection of mini-circle DNA encoding IL-4 was used to overexpress IL-4 from hepatocytes *in vivo* [48]. Serum levels of IL-4 after injection of 0.25 and 0.5 µg IL-4-encoding mini-circle DNA ranged between 1 and 2 ng/ml, whereas IL-4 was not detectable in control mice (Fig. 3A). Peritoneal lavage cells were obtained 24 hours after i.p. infection with 4 × 10^7^ CFU BCG and MINCLE expression on Ly6C^hi^ monocytes and Ly6G^+^ neutrophils was analysed by flow cytometry (Fig. 3B, gating strategy). Compared to the PBS control mice, MINCLE cell surface expression increased after BCG-infection in monocytes and in neutrophils, which was abrogated by IL-4 overexpression specifically in monocytes but not neutrophils (Fig. 3C).

**Figure 3:**
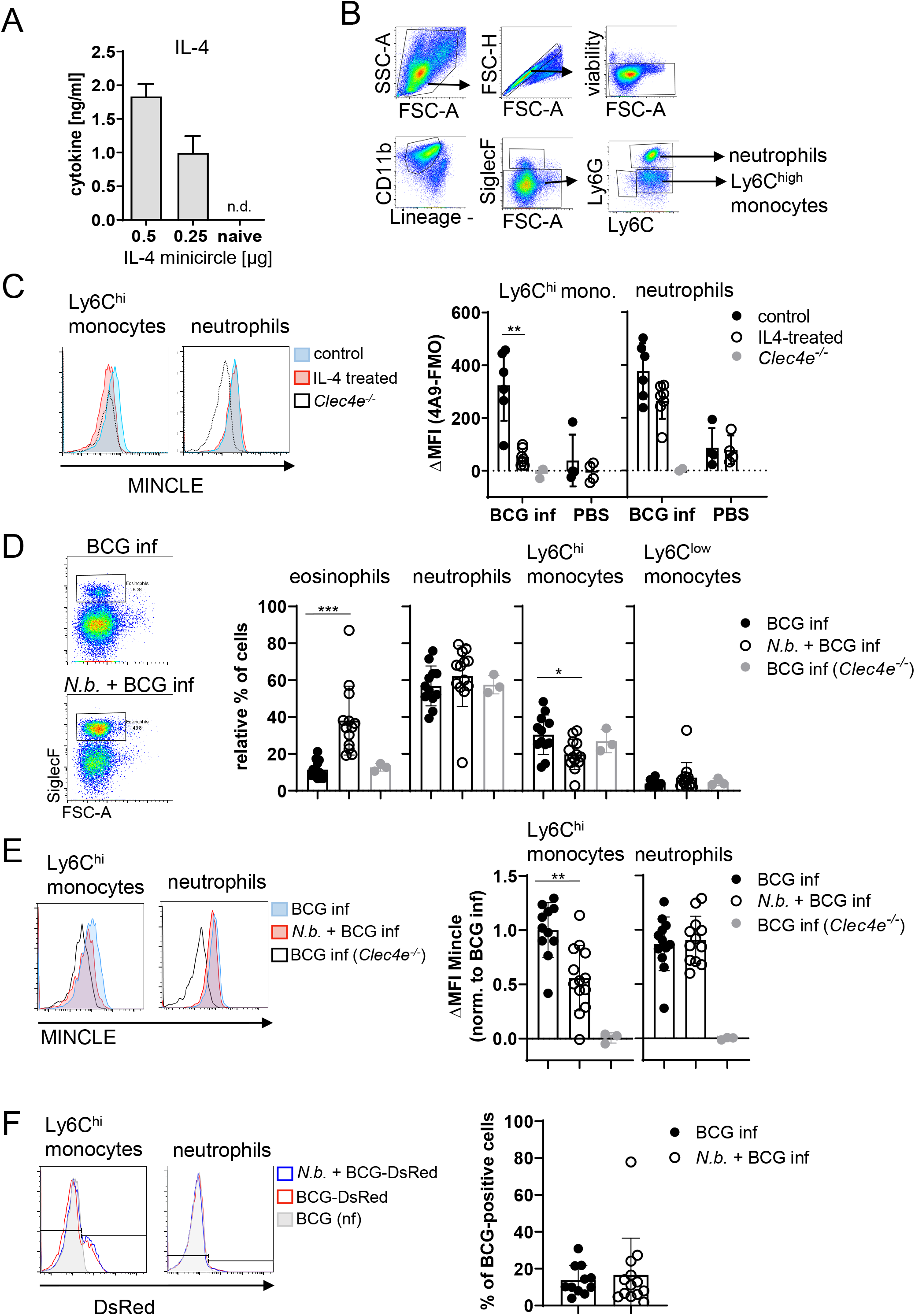
Overexpression of IL-4 or co-infection with *N. brasiliensis* impair MINCLE upregulation on peritoneal monocytes but does not reduce phagocytosis upon BCG infection. **(A)** IL-4 concentration in serum of mice injected with IL-4 minicircle. C57BL/6 mice were hydrodynamically injected with 0.25 µg or 0.5 µg of IL-4 plasmid (i.v.) or Ringer solution. 5 days (0.5 µg) or 7 days later (0.25 µg) mice were sacrificed and serum IL-4 levels were determined via ELISA (n = 6-7 mice per group). n.d. = not detectable. **(B, C)** 0.25 µg of IL-4 plasmid or Ringer solution was hydrodynamically injected into C57BL/6 wildtype mice. 2 days later mice were infected i.p. with 40×10^6^ CFU of *M. bovis* BCG. **(B)** Generic gating strategy for flow cytometry data. Monocytes were characterized as lineage^-^CD11b^+^SiglecF^-^Ly6C^+^ cells. Neutrophils were characterized as lineage^-^CD11b^+^SiglecF^-^Ly6G^+^ cells. Lineage marker: CD3, CD19, NK1.1. **(C)** Histograms depict MINCLE surface expression on Ly6C^hi^ monocytes and neutrophils 24h p.i. analysed via flow cytometry. Quantitative analysis of MINCLE surface expression shown as median fluorescence intensity (MFI). Fluorescence minus one control (FMO) was substracted. Infected *Clec4e*^*-/-*^mice were used as staining controls to exclude unspecific binding of 4A9 antibody. Data is depicted from 2 independent experiments (2-7 mice per group in total, each dot corresponding to one mouse). **p < 0.01; ns = not significant. **(D-E)** C57BL/6 mice were s.c. infected with *N. brasiliensis* or left uninfected followed by *M. bovis* BCG-Dsred infection (40×10^6^ CFU/mouse) on day 10 p.i. 24 h later mice were sacrificed. **(D)** Representative histograms of eosinophil population of BCG-infected and *N. brasiliensis* co-infected mouse. Bar graphs show relative percentage of myeloid cell populations as indicated. **(E)** MINCLE surface expression was analyzed on peritoneal Ly6C^hi^ monocytes and neutrophils via flow cytometry. Quantitative analysis of MINCLE surface expression shown as median fluorescence intensity (MFI) normalized to BCG infected mice. Fluorescence minus one control (FMO) was substracted. Data is depicted from 3 independent experiments (12-13 mice per group in total). *p < 0.05, ** p < 0.01, ***p < 0.001. **(F)** C57BL/6 mice were s.c. infected with *N. brasiliensis* followed by *M. bovis* BCG-Dsred infection (40×10^6^ CFU/mouse) or non-fluorescent BCG (nf) on day 10 p.i. 24 h after BCG infection, phagocytosis was measured by detection of PE signal in monocytes or neutrophils. Quantitative analysis of the percentage of BCG-positive monocytes.

### Co-infection with Nippostrongylus brasiliensis impairs MINCLE upregulation on peritoneal monocytes, but does not reduce phagocytosis, upon BCG infection

Infection with the hookworm *N. brasiliensis* induces a strong Th2 response characterized by high levels of IL-4. We therefore asked whether pre-existing infection with *N. brasiliensis* interferes with MINCLE expression on myeloid cells during mycobacterial infection. The peritoneal cavity of *N. brasiliensis*-infected mice contained a much higher proportion of SiglecF^+^ eosinophils (nearly 40% compared to 10% in controls), whereas the fraction of Ly6C^hi^ monocytes was reduced (from 30% to 20%), and the low percentage of resident monocytes/macrophages was not altered by helminth infection (Fig. 3D). Eosinophils were negative for MINCLE surface staining. MINCLE staining on inflammatory monocytes from BCG-infected mice was significantly reduced when co-infected with *N. brasiliensis* compared to control mice infected only with BCG (Fig. 3E). In contrast, the cell surface expression of MINCLE on neutrophils was not altered by underlying *N. brasiliensis* infection (Fig. 3E). The use of fluorescent BCG-DsRed enabled us to determine the cell types and percentages of peritoneal cells that had ingested mycobacteria 24 hours after injection (Fig. 3F). While on average 12% of Ly6C^hi^ monocytes contained BCG-DsRed, no specific signal was measurable for the peritoneal neutrophils. Co-infection with *N. brasiliensis* did not change the phagocytosis of BCG by inflammatory monocytes in the peritoneal cavity (Fig. 3F).

### Co-infection with N. brasiliensis or with Schistosoma mansoni suppresses Th1/Th17 induction by a MINCLE-dependent adjuvant in the spleen

Downregulation of MINCLE expression on monocytes of mice with *N. brasiliensis* infection led us to ask whether vaccination responses to protein antigen induced by a MINCLE-dependent adjuvant would be inhibited by helminth infections. We investigated this question in chronic (non-transient) *S. mansoni* and acute (transient) *N. brasiliensis* infection models. Infected and control mice were immunized with the recombinant fusion protein H1, comprising the MTB antigens Ag85B and ESAT-6, adsorbed to the adjuvant CAF01 (a combination of the MINCLE ligand TDB incorporated into cationic liposomes) [49]. Subcutaneous injection of H1/CAF01 is known to induce robust IFNγ and IL-17 production by T cells within seven days that is dependent on MINCLE-FcRγ as well as on MyD88 signalling [50].

*S. mansoni* causes a chronic infection in mice, with fertile, adult worms residing in the portal vein system. Trapping of eggs in tissues (especially the liver) is a strong stimulus for Th2-type immunity, which dominates in the later phase of infection after 8-9 weeks. We therefore immunized mice infected with *S. mansoni* in this chronic phase subcutaneously with H1/CAF01 in the flank of the mice (Fig. 4A). We first analysed whether schistosomal infection indeed enhanced IL-4 production by T cells (Fig. 4B). While only very low levels of IL-4 were detectable in the supernatants of draining lymph node cells, splenocytes of *S. mansoni*-infected mice produced significant amounts of IL-4 when stimulated with anti-CD3 *in vitro*. Thus, *S. mansoni*-infection generated a Th2 milieu in the spleen, but not in the inguinal lymph node. Induction of antigen-specific Th1 and Th17 cells, detected by robust secretion of IFNγ and IL-17, respectively, was observed after immunization in both lymph nodes and spleen (Fig. 4C and D). Infection with *S. mansoni* had not impact on H1-specific or anti-CD3-induced production of IFNγ, IL-17 and of IL-10 from draining inguinal lymph nodes (Fig. 4C). In contrast, the antigen-specific splenocyte responses to H1 were strongly reduced in the case of IFNγ, IL-17 and also for IL-10 (Fig. 4D). *S. mansoni* infection also suppressed IFNγ, but not IL-17 production triggered by polyclonal anti-CD3 stimulation, whereas secretion of IL-10 was much higher from splenocytes of *S. mansoni*-infected mice (Fig. 4D). Together, these data show that *S. mansoni*-infection establishes a Th2 environment in the spleen but not in the inguinal lymph nodes, which corresponds to suppression of Th1 and Th17 responses after CAF01-adjuvanted immunization in splenocytes but has no impact on draining lymph node cells.

**Figure 4:**
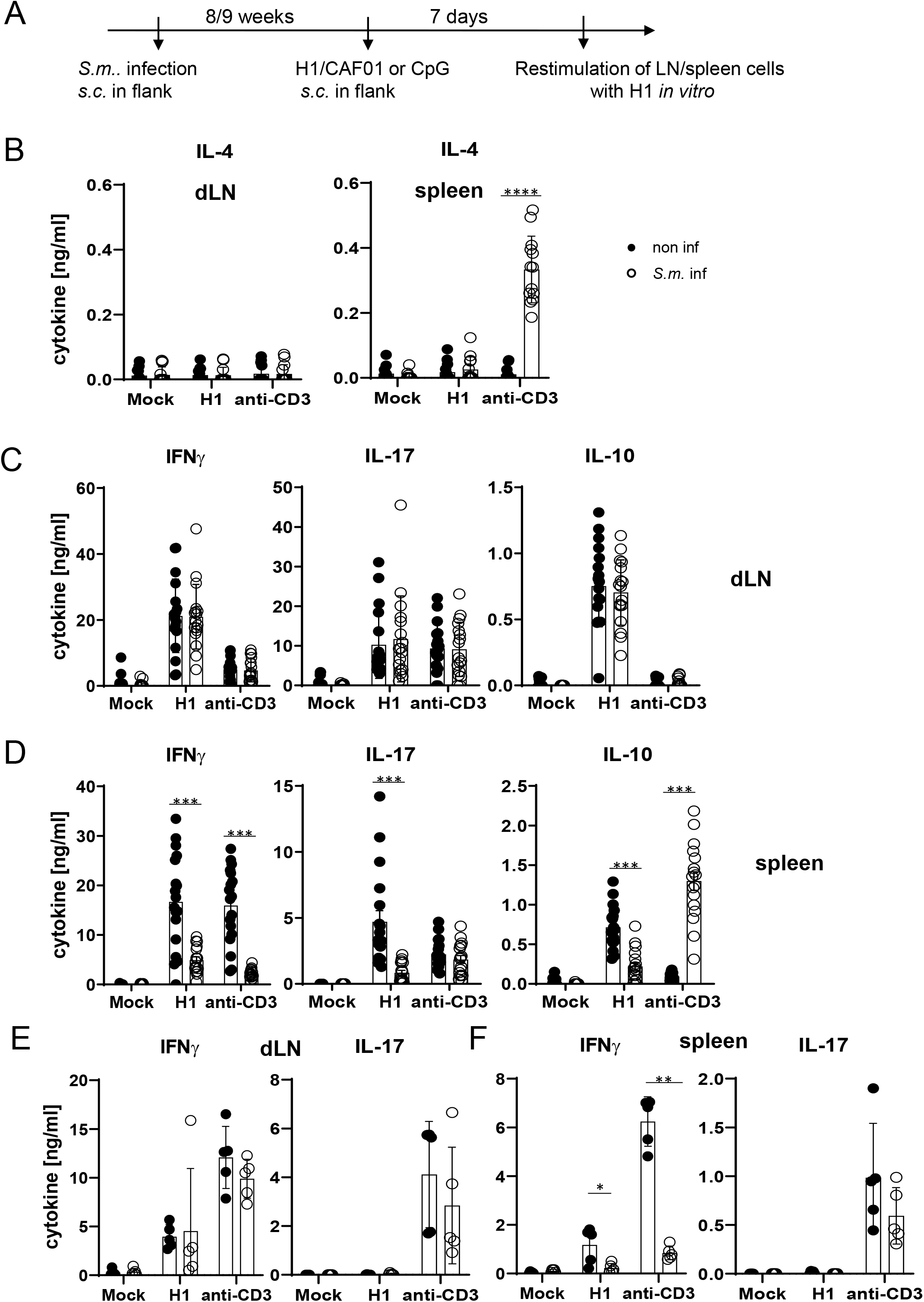
Co-infection with *Schistosoma mansoni (S*.*m*.) suppresses Th1/Th17 induction by a MINCLE-dependent adjuvant in the spleen but not in the draining lymph node. **(A)** Scheme of experimental procedure. *Cercariae* of *Schistosoma mansoni* (*S*.*m*.) were injected s.c into C57BL/6 mice. 8 to 9 weeks p.i. mice were immunized with CAF01 **(B, C, D)** or CpG ODN **(E, F)**. 7 days after immunization, mice were sacrificed and inguinal lymph node cells and spleen cells were re-stimulated with H1 *in vitro*. Draining inguinal lymph node cells or splenocytes were restimulated with H1 or anti-CD3 for 96 h. IL-17, IFNγ, IL-10 and IL-4 production was measured from cell culture supernatants by ELISA. Data is shown from 3 independent experiments (n = 18 mice per group in total) for IFNγ, IL-17 and IL-10 **(C, D)** and from 2 independent experiments for IL-4 **(B). (E, F)** Data is shown from 1 experiment (n = 5 mice per group in total). *p < 0.05, **p < 0.01, ***p < 0.001.

To determine whether *S. mansoni*-induced inhibition of antigen-specific immune responses by splenocytes, but not draining lymph node cells, was specific for the MINCLE-dependent adjuvant CAF01, we immunized mice with H1 together with the TLR9 ligand CpG ODN 1826 as adjuvant. Using the same protocol as before for H1/CAF01, we observed, as demonstrated previously [28], strong induction of IFNγ-producing T cells but a lack of IL-17 production upon restimulation of draining lymph node cells or splenocytes (Fig. 4E, F). Thus, it was not possible to compare inhibition of Th17 responses by helminth infection between adjuvants. However, *S. mansoni* did not generally down-regulate IL-17 production from T cells, as the levels after anti-CD3-stimulation were unaffected (Fig. 4E, F). Similar to the impact on immunization with CAF01, *S. mansoni* infection strongly reduced antigen-specific and non-specific IFNγ secretion by splenocytes but not by draining lymph node cells (Fig. 4E, F). Thus, *S. mansoni* infection caused a general suppression of the splenic Th1 response to immunization, regardless whether the MINCLE-dependent adjuvant CAF01 was used or the TLR9-dependent CpG ODN 1826.

To address the effect of transient helminth infection, mice were injected subcutaneously with L3 larvae of *N. brasiliensis* and were immunized five days later with H1/CAF01 in the footpads of both hindlegs (Fig. 5A). The local tissue swelling to subcutaneous vaccination in the footpad was not changed in mice with underlying *N. brasiliensis* infection (Fig. 5B). The total number of cells in the draining inguinal lymph nodes 7 days after immunization was strongly increased compared to non-vaccinated mice, but significantly reduced in mice infected with *N. brasiliensis*; in contrast, the number of splenocytes was unaltered by helminth infection (Fig. 5C). As was already observed after infection with *S. mansoni*, establishment of IL-4-producing Th2 T cells was only detected in spleens of *N. brasiliensis*-infected mice but not in their popliteal and inguinal lymph nodes (Fig. 5D). Upon restimulation of lymph node cells with H1 *in vitro*, the antigen-specific production of IFNγ, IL-17 or IL-10 was not affected by *N. brasiliensis* co-infection (Fig. 5E). In striking contrast, splenocytes from co-infected mice generated significantly less IFNγ and IL-17, but not IL-10, when restimulated with H1 antigen, whereas for polyclonal stimulation with anti-CD3 antibody only a reduction for IFNγ was observed (Fig. 5F). When mice were immunized with H1 together with the TLR4-dependent adjuvant G3D6A, IFNγ production by splenocytes was diminished in mice infected with *N. brasiliensis*, whereas antigen-specific secretion of IL-17, IL-10 and IL-4 were not changed (Fig. 5G). However, polyclonal stimulation of T cells from the spleens of N. brasiliensis-infected mice caused secretion of significant IL-4 (Fig. 5G), as after immunization with the MINCLE-dependent adjuvant CAF01 (Fig. 5D).

**Figure 5:**
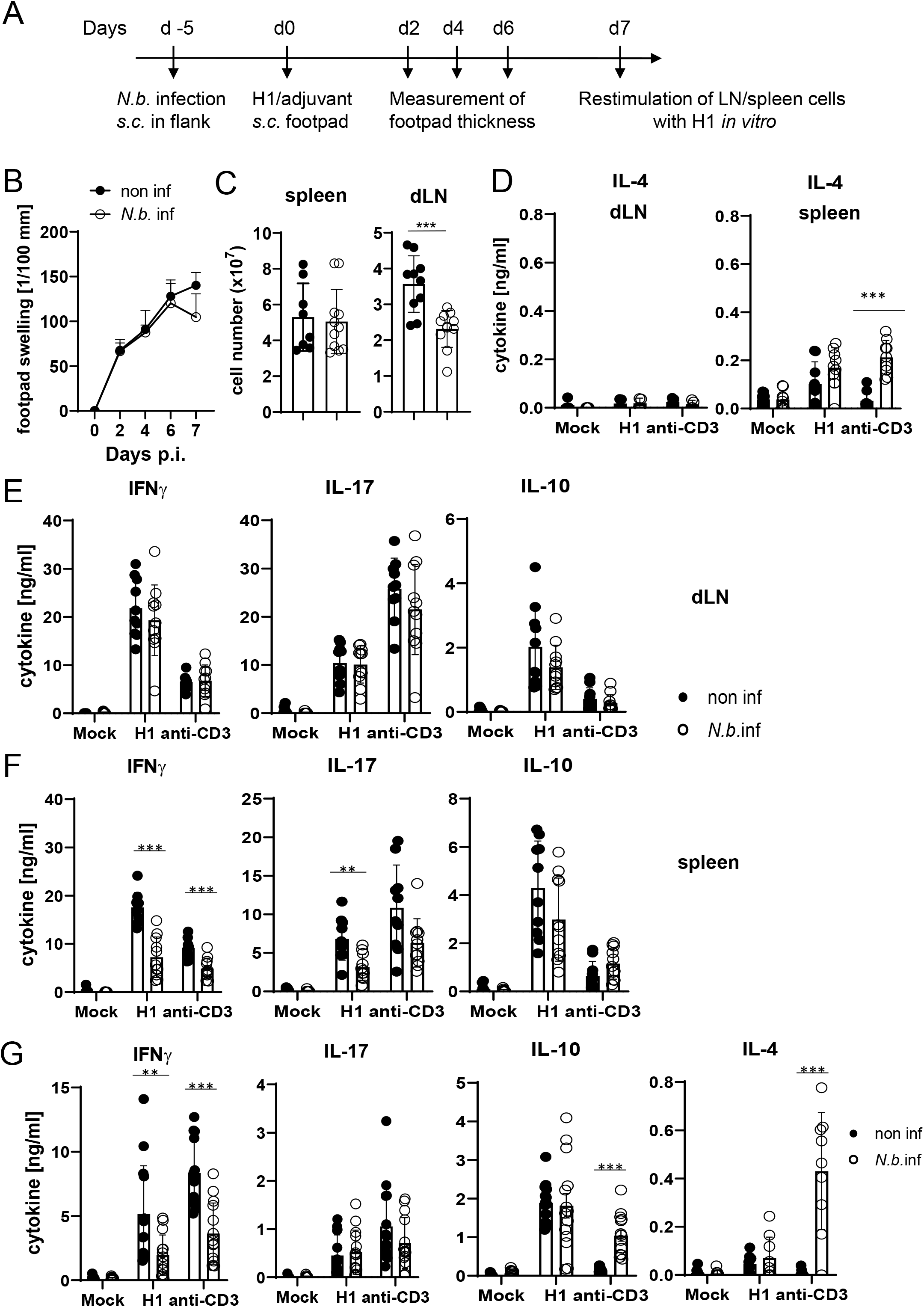
Co-infection with *N. brasiliensis* suppresses Th1/Th17 induction by a MINCLE-dependent adjuvant in the spleen but not in the draining lymph node. **(A)** Scheme of experimental procedure. C57BL/6 mice were infected subcutaneously in the flank with 500 L3 *larvae* of *N. brasiliensis* in 200 µl PBS. 5 days p.i. mice were immunized with H1/CAF01 **(B-F)** or H1/G3D6A **(G)**. 7 days after immunization, mice were sacrificed, and inguinal and popliteal lymph node cells and spleen cells were re-stimulated with H1 *in vitro*. **(B)** Footpad swelling was measured over a period of 1 week after immunization with H1/CAF01 until mice were sacrificed (day 7 after immunization). The increase in footpad swelling is shown as mean + SD for the indicated time points (n = 10-11 mice for each data point). **(C)** Absolute cell number of draining inguinal and popliteal lymph node cells (LN) and spleen on day 7 after H1/CAF01. **(D-G)** Draining inguinal and popliteal lymph node cells or splenocytes were restimulated with H1, anti-CD3 or left untreated (mock) for 96 h, followed by cytokine determination by ELISA. **(D)** IL-4 levels produced by draining lymph node cells (left) or splenocytes (right). **(E, F)** IL-17, IFNγ and IL-10 cytokine production by draining lymph node cells **(E)** and splenocytes **(F). (G)** Immunization of C57BL/6 mice with H1 in G3D6A adjuvant. 7 days after immunization mice were sacrificed and splenocytes were treated as described in **(D, E)**. All data is shown from two **(B-F)** or three **(G)** independent experiments (n = 10-14 mice per group in total). p < 0.05, ** p < 0.01, *** p < 0.001.

Together, these results indicate that underlying *N. brasiliensis* infection attenuated cell expansion in the draining lymph node, yet the differentiation toward Th1/Th17 at this site was not affected. In contrast, hookworm infection did interfere with antigen-specific IFNγ and IL-17 production by splenic T cells, correlating with strongly enhanced production of IL-4 by splenocytes (but not draining lymph node cells). The inhibitory effect of infection on IFNγ appears to be general, antigen- and adjuvant-non-specific, because it was also observed after polyclonal T cell stimulation and independent of the pattern recognition receptor pathway triggered by the adjuvant used. In contrast, the impairment of Th17 induction in the spleen by *N. brasiliensis* was specific for the MINCLE-dependent CAF01. Taken together, immunization experiments in two helminth infection models demonstrated organ-specific inhibition of Th responses in the spleen, with antigen-specific inhibition of CAF01-induced, MINCLE-dependent IL-17 production.

## Discussion

In this study, we show for the first time that IL-4 impairs MINCLE expression and function *in vivo*. Two different models of helminth infection revealed an organ-specific impairment of Th17 induction by the MINCLE-dependent adjuvant CAF01 in the spleen. These findings establish an *in vivo* impact of IL-4 on the expression and function of MINCLE that may have important consequences for the detection of and the response to mycobacteria and their cell wall glycolipids in infection or vaccination.

In previous work, we have found that IL-4 downregulates expression of DECTIN-2 family CLRs in murine and human macrophages and DC [43]. Here, we first extended these findings by showing that upregulation of these CLR by BCG was also impaired by IL-4 in BMM. Importantly, using overexpression of IL-4 *in vivo* by hydrodynamic injection of minicircle DNA, the upregulation of MINCLE on monocytes recruited after BCG infection to the peritoneum was strongly inhibited. This effect was cell type-specific, as it was not observed on recruited neutrophils, and it was also caused by infection with *N. brasiliensis*.

IL-4 and *N. brasiliensis* infection did not interfere with phagocytosis of BCG *in vitro* or *in vivo*, showing that high level expression of DECTIN-2 family CLR is not required for phagocytosis of mycobacteria. This may seem surprising, as DECTIN-2 and MCL are phagocytic receptors [51, 52], and mycobacteria strongly express ligands for both receptors (mannans and TDM). A lack of DECTIN-2 family CLR appears to be compensated by the large number of other receptors contributing to phagocytosis of mycobacteria, including the mannose receptor CD206, integrins like CD11b, or the inhibitory CLR DC-SIGN [53-55].

Importantly, IL-4-treated macrophages not only down-regulated DECTIN-2 family CLR, but in addition produced significantly less cytokines after stimulation with BCG. A similar reduction in G-CSF and TNF production was observed here in MINCLE- and FcRγ-deficient BMM. These observations suggests that IL-4 produced during helminth infection could interfere with the initial sensing of invading mycobacteria by pulmonary macrophages and thereby mute the generation of an inflammatory chemokine/cytokine as well as an anti-microbial response. If so, impaired detection of inhaled MTB by alveolar and lung macrophages may favour intracellular mycobacterial survival and replication, providing a mechanism underlying the reported increase in tuberculin skin test conversion reported for helminth-infected household contacts of patients with smear-positive tuberculosis [6]. Whether IL-4 impairs functional macrophage responses to mycobacteria primarily through downregulation of MINCLE and DECTIN-2 family CLRs or by inhibiting other cytokine-inducing pathways is an open question that needs to be investigated in future work.

In addition to the response of macrophages to BCG, we investigated how type 2 immune bias during helminth infections impacts on the Th cell differentiation after recombinant subunit vaccination with MINCLE-dependent and –independent adjuvants. Induction of Th17 immunity triggered by the MINCLE-dependent adjuvant CAF01 was suppressed in two models of helminth infection in the spleen, but not in the draining popliteal and inguinal lymph nodes. This organ-specific impact of infection with *N. brasiliensis* and *S. mansoni* on splenic Th cell differentiation was associated with a stronger Th2 bias in the spleen, as demonstrated by the robustly increased levels of IL-4 and IL-10 released by splenocytes, but not draining lymph node cells, from worm-infected mice after polyclonal stimulation with anti-CD3 (Figs. 4 and 5). Compartmentalized overexpression of IL-10 in the spleens but not lymph nodes has been described before in mice with chronic schistosomiasis [56]. Our data suggest that such a restriction of Th2 bias to the spleen also applies to IL-4 producing T cells in both helminth infections.

Helminth infection did not alter the production of IL-17, IFNγ or IL-10 by antigen-specific CD4+ T cells in the draining lymph nodes, as determined by restimulation of a defined number of cells with recombinant H1. However, the cellularity of the lymph nodes was significantly diminished by *N. brasiliensis*, suggesting that the total number of antigen-specific Th1 / Th17 cells in the draining lymph nodes is reduced by co-existent worm infection. Decreased peripheral lymph node size and cellularity has been reported in mice infected with the enteric helminth *H. polygyrus* and was associated with a dampened immune response to infection with BCG, affecting primarily B and T lymphocytes [57]. In the *H. polygyrus* model, the gut-draining mesenteric lymph nodes were increased in size, suggesting that atrophy of peripheral lymph nodes was due to a redistribution of lymphocytes [57]. Furthermore, in mice infected with *Helicobacter pylori*, we have recently shown that concomitant schistosome infection lead to a redistribution of bacterial-specific CXCR3+ T cells resulting in reduced bacterial growth control [58]. Cell numbers and size of lymph nodes are controlled by entry of lymphocytes *via* the high endothelial venules and exit through the efferent lymphatics.

While the inhibition of IFNγ production by splenocytes from helminth-infected mice was impaired also after immunization with the TLR4 ligand G3D6A and the TLR9 ligand CpG ODN as adjuvants, IL-17 production was specifically impaired when the MINCLE-dependent CAF01 was the adjuvant. In addition, antigen-non-specific production of IFNγ by polyclonal anti-CD3 stimulation was also inhibited in splenocytes from helminth-infected mice, suggesting a general inhibitory effect of *N. brasiliensis* and *S. mansoni* infection that was not observed for production of IL-17. Thus, underlying helminth infection affects the CAF01-induced antigen-specific Th cell response in the spleen in a manner that could be explained by down-regulation of MINCLE expression. Whether this is indeed the case needs to be further investigated in future studies comparing myeloid cells in spleen, peripheral lymph node and subcutaneous injection site tissue. Detection and quantification of MINCLE surface protein on myeloid cells from tissues by flow cytometry is unfortunately complicated by the fact that proteolytic agents commonly used for enzymatic tissue-dissociation cause a near complete loss of specific staining [59]. For this reason, we employed here the intraperitoneal BCG-infection model for assessment of IL-4/helminth effects on myeloid cells *in vivo*, because it allowed the direct flow cytometric staining of peritoneal exudate cells without the need for enzymatic tissue digestion.

Interestingly, application of schistosome eggs, a strong inducer of IL-4 and IL-13 production *in vivo*, prior to infection of mice with *Salmonella typhimurium* downregulated the Th17 response in the gut mucosa and impaired the clearance of the bacteria from the gut [60]. As the *Salmonella* cell wall contains trehalose phospholipids that bind to and activate MINCLE [61], it is possible that induction of IL-17-producing cells during infection requires MINCLE and may be inhibited by schistosome eggs through IL-4/IL-13-induced downregulation of the receptor. Alternatively, IL-4 and IL-10 present in the spleen during helminth infection may act directly on antigen-specific Th cell differentiation after immunization and block Th17 [62, 63] and Th1 bias [64].

More generally, down-regulation of other PRR in addition to DECTIN-2 family CLR during acute or chronic helminth infection may also have a negative impact on vaccines comprising specific PAMPs as adjuvants. Indeed, several papers have demonstrated a diminished baseline expression of TLR2 and TLR9 patients with filarial infection [65, 66]. Underlying infection with hookworm or filaria were found to down-regulate Th1 and Th17 responses in individuals with latent tuberculosis [67, 68]. Certainly, helminth-induced immune regulation can interfere by diverse and manifold mechanisms with Th1/Th17 adaptive immunity. It will be worthwhile to investigate further, in animal models and in human patients with helminth infection, whether impaired DECTIN-2 family CLR expression is a significant contributor to thwarting of these protective anti-mycobacterial T cell responses.

## Methods

### Mice

C57BL/6 wildtype, MINCLE knockout (*Clec4e*^*-/-*^) and FcRγ knockout (*Fcerg1*^-/-^) mice were bred under specific pathogen-free (SPF) conditions at the “Präklinische Experimentelle Tierzentrum” (PETZ) of the Medical Faculty in Erlangen. *Clec4e*^*-/-*^ mice were generated by the Consortium for Functional Glycomics [69] and used with permission. *Fcerg1*^*-/-*^ mice [70] were kindly provided by Dr. Falk Nimmerjahn. C57BL/6N mice were purchased from Charles River Laboratories. All mouse experiments were approved by the “Regierung von Unterfranken” (protocol number 55.2.2-2532-543) and the “Regierung of Oberbayern” (protocol number ROB-55.2 Vet_02-17-145). Male mice between 8 and 12 weeks of age were used for *in vivo* experiments.

### Bacteria

*Mycobacterium bovis* (BCG) was grown in Middlebrook 7H9 broth supplemented with 10 % OADC-enrichment medium and 0.05 % Tween 80 in small cell culture flasks constantly shaking at 125 rpm at 37 °C to an OD_600_ ∼ 1-2. Prior to *in vitro* stimulation, BCG were washed with PBS and diluted in cDMEM.

### Hydrodynamic injection of IL-4 minicircle DNA

To investigate IL-4 derived effects on immunization responses or BCG-infections in mice, we utilized a technique that leads to a systemic overexpression of IL-4. A minicircle DNA vector that encodes the gene for IL-4, but lacks further bacterial elements in comparison to conventional plasmids, was hydrodynamically injected intravenously in the tail vein of five-week old male mice [71]. 0.5 µg of IL-4 was injected in 2 ml Ringer solution in under 10 seconds per mouse.

### Helminth infections

All *Schistosoma mansoni* infection experiments were performed at TU Munich. C57BL/6 mice were infected subcutaneously as described previously [58] with 100 cercariae from the NMRI strain (originally from Puerto Rico) of *S. mansoni* in 100 µl PBS shed by infected *Biomphalaria glabrata* snails, provided by the NIAID Schistosomiasis Resource Center of Biomedical Research Institute (Rockville, MD) through NIH-NIAID Contract HHSN272201700014I for distribution through BEI Resources. Mice were vaccinated after 8 weeks of infection, when parasite egg-induced Th2 immune responses begin to peak.

The *Nippostrongylus brasiliensis* life cycle was maintained in rats. *N. brasiliensis* larvae were cultured in a mixture of charcoal and feces of infected rats in petri dishes for 15-40 days at 37°C. Subsequently, third-stage (L3) larvae were harvested with 10 ml of 0.9 % saline and transferred to a Baermann apparatus. Within 1h larvae descended to the bottom of the funnel. Larvae were then transferred to a fresh 50 ml tube and extensively washed by repeating steps of sedimentation and exchange of saline. C57BL/6 mice were infected subcutaneously with 500 L3 larvae in 200 µl PBS in the flank of each mouse using a 25 G needle. All worms were expelled completely within 10 days.

### BCG infection of mice

10 days after *N. brasiliensis* infection, or 2 days after hydrodynamic injection of IL-4 minicircle DNA, mice were infected intraperitoneally with 40×10^6^ CFU of *M. bovis* BCG in a volume of 200 µl PBS using a 27-G needle. PBS-injected mice as well as completely naïve mice were used as controls. Mice were sacrificed 24 h post infection.

### Immunizations

Mice were immunized subcutaneously with 50 µl of a mixture of 1 µg H1, a fusion protein of the MTB antigens Ag85B and ESAT-6, and the respective adjuvants (CAF01, CpG ODN 1826, or G3D6A) in the footpads of the hind legs or in the flank as indicated in the Figure legends. CAF01 is composed of TDB and cationic dimethyldioctadecylammonium (DDA) liposomes and has been described in detail before [49]. CpG ODN 1826 is a phosphorothioate-protected oligonucleotide and was synthetized by TIB MOLBIOL (Berlin, Germany). G3D6A is a liposomal adjuvant formulation. It is comprised of the synthetic TLR4 ligand 3-O-de-acyl-hexaacyl-monophoshoryl lipid A (3D-6A-SMPLA, 3D(6-acyl)-PHAD®) manufactured as cGMP product by Avanti Polar Lipids Inc. (Alabaster, Alabama,U.S.A) and embedded in a matrix of 1,2-dimyristoyl-sn-glycero-3-phosphocholine (DMPC), 1,2-Dimyristoyl-sn-glycero-3-phosphoglycerol (DMPG) and Cholesterol (Molar ration: 9:1:7.5) in an aqueous suspension buffered with PBS. Footpad swelling was monitored regularly and measured prior to immunization as well as every second day post immunization. On day 7 post immunization, inguinal and popliteal lymph nodes were analysed.

### Isolation and culture of bone marrow-derived macrophages (BMM)

Bone marrow cells were differentiated to BMM for 6-7 days in complete Dulbecco’s Modified Eagle Medium containing 10 % FCS, 50 μM β-mercaptoethanol and Penicillin/Streptomycin (cDMEM) supplemented with 10 % L929-cell conditioned medium (LCCM) as a source of M-CSF.

### Stimulation of BMM

BMM were stimulated with plate-coated TDB (Polar Avanti, 5µg/ml), using isopropanol as a mock control, as described [28], LPS (*E. coli* serotype O55:B5, Sigma, 10ng/ml), CpG ODN 1826 (TIB MOLBIOL, 0.5 µM), or with BCG at the indicated MOI.

### Restimulation of lymph node cells

Inguinal and popliteal lymph nodes were collected and meshed through a 70 µm nylon filter to get a single cell suspension. 5×10^5^ cells were stimulated in 96-well U-bottom plates with H1 (1 µg/ml), soluble anti-CD3 (0.5 µg/ml) or left untreated (mock) for 96 h.

### mRNA expression of CLR

RNA was isolated using Trifast (Peqlab) and transcribed to cDNA (High capacity cDNA Reverse Transcription Kit, Applied Biosystems). Expression of the house keeping gene *Hprt* and of the genes of interest was analyzed by quantitative real time PCR (qRT-PCR). All primers and probes were selected from the Roche Universal Probe Library (UPL). Ct values of the target genes were normalized to *Hprt*, calibrated to unstimulated cells, and depicted as fold change.

### Cytokine ELISA

Secreted cytokines were analyzed by ELISA (R&D Systems) from cell culture supernatants of stimulated BMM or lymph node cells.

### Flow cytometry

Cell surface expression of MINCLE was analyzed by flow cytometry. Cells were blocked with anti-mouse CD16/32, stained with the primary antibody anti-MINCLE (clone 4A9, MBL, 1µg/ml), followed by anti-rat IgG1-APC (eBioscience). FACS data were acquired on a LSRFortessa™ (BD) and analyzed using the software FlowJo (v10).

### Statistics

GraphPad Prism software (version 8) was used for statistical analysis. Statistical significance was calculated using Mann Whitney U-test to compare two non-paired groups. * p < 0.05, ** p < 0.01, *** p < 0.001, *ns* p > 0.05.

## Acknowledgements

Animal husbandry by Manfred Kirsch, technical assistance by Nina Grohmann, and support by Christian Bogdan is gratefully acknowledged. We thank Nathalie Thuma and Paul Haase for help with *N. brasiliensis* infections.

This work was funded by Deutsche Forschungsgemeinschaft (GRK 1660-TP-A02 and LA 1262/8-1 to R.L., and CRC1181_A02 to D.V., and CO 1469/16-1 to C.P.d.C.).

## Figure legends

**Figure 1: IL-4 impairs upregulation of MINCLE and other DECTIN-2 family CLR in macrophages stimulated with BCG**. (A-D) C57BL/6 BMM were stimulated as indicated in presents or absence of IL-4 for 6, 24 or 48h. MINCLE (A), DECTIN-2 (B), MCL (C), and DECTIN-1 (D) mRNA expression was determined by qRT-PCR shown as fold change calibrated to unstimulated control. Data are depicted as mean + SD from two independent experiments performed in biological duplicates. (E) BMM were stimulated with BCG at the indicated MOI in the presence or absence of IL-4 for 24 hours, followed by staining of MINCLE expression. Representative stainings are shown as histogram overlay (left panel), for quantification (right panel) the median fluorescence intensity of the isotype control staining was subtracted from MINCLE signal to obtain the ΔMFI. Each point represents one mouse, pooled from two experiments. *p < 0.05, **p < 0.01, ***p < 0.001.

**Figure 2: IL-4 does not affect phagocytosis of BCG but inhibits cytokine production. (A)** C57BL/6 BMM were infected with different MOI of fluorescent BCG-Dsred co-treated with IL-4 or not as indicated. Phagocytic uptake was measured via flow cytometry. **(A)** Representative histograms show phagocytosis of BCG by detection of fluorescent DsRed signal in BMMs. Quantitative analysis of phagocytosis based on percentage of BCG-positive cells at 6 and 24 h post infection. Data is depicted from 2-3 independent experiments performed in biological duplicates. **(B)** C57BL/6 *(WT)*, MINCLE knockout *(Clec4e*^*-/-*^) and FcRγ (*Fcerg1*^*-/-*^) BMM were stimulated with BCG for 24h. Production of G-CSF and TNF was measured from cell culture supernatants via ELISA. No significant cytokine production was detected from non-stimulated BMM. Data is depicted from 3 independent experiments performed in biological duplicates. *p < 0.05, **p < 0.01, ***p < 0.001.

**Figure 3: Overexpression of IL-4 or co-infection with *N. brasiliensis* impair MINCLE upregulation on peritoneal monocytes but does not reduce phagocytosis upon BCG infection. (A)** IL-4 concentration in serum of mice injected with IL-4 minicircle. C57BL/6 mice were hydrodynamically injected with 0.25 µg or 0.5 µg of IL-4 plasmid (i.v.) or Ringer solution. 5 days (0.5 µg) or 7 days later (0.25 µg) mice were sacrificed and serum IL-4 levels were determined via ELISA (n = 6-7 mice per group). n.d. = not detectable. **(B, C)** 0.25 µg of IL-4 plasmid or Ringer solution was hydrodynamically injected into C57BL/6 wildtype mice. 2 days later mice were infected i.p. with 40×10^6^ CFU of *M. bovis* BCG. **(B)** Generic gating strategy for flow cytometry data. Monocytes were characterized as lineage^-^CD11b^+^SiglecF^-^Ly6C^+^ cells. Neutrophils were characterized as lineage^-^CD11b^+^SiglecF^-^Ly6G^+^ cells. Lineage marker: CD3, CD19, NK1.1. **(C)** Histograms depict MINCLE surface expression on Ly6C^hi^ monocytes and neutrophils 24h p.i. analysed via flow cytometry. Quantitative analysis of MINCLE surface expression shown as median fluorescence intensity (MFI). Fluorescence minus one control (FMO) was substracted. Infected *MINCLE-/-* mice were used as staining controls to exclude unspecific binding of 4A9 antibody. Data is depicted from 2 independent experiments (2-7 mice per group in total, each dot corresponding to one mouse). **p < 0.01; ns = not significant. **(D-E)** C57BL/6 mice were s.c. infected with *N. brasiliensis* or left uninfected followed by *M. bovis* BCG-Dsred infection (40×10^6 CFU/mouse) on day 10 p.i. 24 h later mice were sacrificed. **(D)** Representative histograms of eosinophil population of BCG-infected and *N. brasiliensis* co-infected mouse. Bar graphs show relative percentage of myeloid cell populations as indicated. **(E)** MINCLE surface expression was analyzed on peritoneal Ly6C^hi^ monocytes and neutrophils via flow cytometry. Quantitative analysis of MINCLE surface expression shown as median fluorescence intensity (MFI) normalized to BCG infected mice. Fluorescence minus one control (FMO) was substracted. Data is depicted from 3 independent experiments (12-13 mice per group in total). *p < 0.05, ** p < 0.01, ***p < 0.001. **(F)** C57BL/6 mice were s.c. infected with *N. brasiliensis* followed by *M. bovis* BCG-Dsred infection (40×10^6^ CFU/mouse) or non-fluorescent BCG (nf) on day 10 p.i. 24 h after BCG infection, phagocytosis was measured by detection of PE signal in monocytes or neutrophils. Quantitative analysis of the percentage of BCG-positive monocytes.

**Figure 4: Co-infection with *Schistosoma mansoni (S.m.)* suppresses Th1/Th17 induction by a MINCLE-dependent adjuvant in the spleen but not in the draining lymph node. (A)** Scheme of experimental procedure. *Cercariae* of *Schistosoma mansoni* (*S*.*m*.) were injected s.c into C57BL/6 mice. 8 to 9 weeks p.i. mice were immunized with CAF01 **(B, C, D)** or CpG ODN **(E, F)**. 7 days after immunization, mice were sacrificed and inguinal lymph node cells and spleen cells were re-stimulated with H1 *in vitro*. Draining inguinal lymph node cells or splenocytes were restimulated with H1 or anti-CD3 for 96 h. IL-17, IFNγ, IL-10 and IL-4 production was measured from cell culture supernatants by ELISA. Data is shown from 3 independent experiments (n = 18 mice per group in total) for IFNγ, IL-17 and IL-10 **(C, D)** and from 2 independent experiments for IL-4 **(B). (E, F)** Data is shown from 1 experiment (n = 5 mice per group in total). *p < 0.05, **p < 0.01, ***p < 0.001.

**Figure 5: Co-infection with *N. brasiliensis* suppresses Th1/Th17 induction by a MINCLE-dependent adjuvant in the spleen but not in the draining lymph node. (A)** Scheme of experimental procedure. C57BL/6 mice were infected subcutaneously in the flank with 500 L3 *larvae* of *N. brasiliensis* in 200 µl PBS. 5 days p.i. mice were immunized with H1/CAF01 **(B-F)** or H1/G3D6A **(G)**. 7 days after immunization, mice were sacrificed, and inguinal and popliteal lymph node cells and spleen cells were re-stimulated with H1 *in vitro*. **(B)** Footpad swelling was measured over a period of 1 week after immunization with H1/CAF01 until mice were sacrificed (day 7 after immunization). The increase in footpad swelling is shown as mean + SD for the indicated time points (n = 10-11 mice for each data point). **(C)** Absolute cell number of draining inguinal and popliteal lymph node cells (LN) and spleen on day 7 after H1/CAF01. **(D-G)** Draining inguinal and popliteal lymph node cells or splenocytes were restimulated with H1, anti-CD3 or left untreated (mock) for 96 h, followed by cytokine determination by ELISA. **(D)** IL-4 levels produced by draining lymph node cells (left) or splenocytes (right). **(E, F)** IL-17, IFNγ and IL-10 cytokine production by draining lymph node cells **(E)** and splenocytes **(F). (G)** Immunization of C57BL/6 mice with H1 in G3D6A adjuvant. 7 days after immunization mice were sacrificed and splenocytes were treated as described in **(D, E)**. All data is shown from two **(B-F)** or three **(G)** independent experiments (n = 10-14 mice per group in total). p < 0.05, ** p < 0.01, *** p < 0.001.

